# Distinct impacts of air and root-zone temperatures on leaf and root features of cucumber seedlings: resource acquisition capacity, organ size and carbon-nitrogen balance

**DOI:** 10.1101/271403

**Authors:** Xiaozhuo Wang, Lihong Gao, Yongqiang Tian

**Author notes:** Correspondence: Lihong Gao Yongqiang Tian. The author responsible for distribution of materials integral to the findings presented in this article in accordance with the policy described in the Instructions for Authors is: Lihong Gao and Yongqiang Tian.

## Abstract

Both low air (*T*_air_) and root-zone (*T*_root_) temperatures can inhibit resource (e.g. carbon and nutrients) acquisition by leaves and roots through various aspects, such as morphology, biomass allocation and assimilation/absorption capacity. However, it is still ambiguous whether *T*_air_ and *T*_root_ influence carbon (C) and nutrient acquisition via the same approach. To this end, in this study, cucumber (*Cucumis sativus* L.) seedlings were hydroponically grown under treatments arranged in complete factorial combination of two levels of *T*_air_ (26/18°C and 20/12°C, day/night) and two levels of *T*_root_ (19°C and 13°C, constant). In general, both *T*_air_ and *T*_root_ affected leaf and root sizes mainly by regulating their morphology rather than biomass investment. Under low *T*_air_ conditions (20/18°C), elevated *T*_root_ (compare 19°C versus 13°C) did not influence C acquisition, but increased nitrogen (N) acquisition mainly due to an increase in relative root length, resulting in decreased C : N acquisition ratio. However, under low *T*_root_ conditions (13°C), elevated *T*_air_ (compare 26/18°C versus 20/12°C) enhanced both C and N acquisition mainly because of an increase of both C assimilation in leaves and N absorption by roots, resulting in relatively constant C : N acquisition ratio. In addition, the *T*_air_ and *T*_root_ interaction was mainly observed in relative growth rate and root growth-related variables. Our results infer that *T*_air_ and *T*_root_ have distinct impacts on resource acquisition and carbon-nitrogen balance in plants.

## INTRODUCTION

Low temperature stress is a commonly encountered problem for plants in most temperate or high-altitude regions during cool-season cultivation. Low temperature may inhibit plant growth in a complex manner. Primarily, it limits the size of leaves and roots per unit plant biomass (leaf area ratio, LAR, and root length ratio, RLR) as an integrated result of altered biomass fraction (leaf mass fraction, LMF, and root mass fraction, RMF) and morphological characteristics (leaf area per unit leaf mass, i.e. specific leaf area, SLA, and root length per unit root mass, i.e. specific root length, SRL) (Tachibana, 1982; Weih and Karlsson, 2001). In addition, it decreases the capacity of resource acquisition per unit size of leaves and roots (Clarkson et al., 1986; Delucia et al., 1992). These two aspects together inhibit the access to resources, and thus retard the relative growth rate (RGR) of plants (Loveys et al., 2002).

Knowledge about the relative contributions of various plant components to RGR may help us better predict plant responses to environmental variation and then pursue the right temperature control strategy. Previous researches (Loveys et al., 2002; Poorter et al., 2012) suggest that, for the above ground part of plants, SLA usually plays a more flexible and important role than LMF in determining LAR, while net assimilation rate (NAR, increase in plant mass per unit leaf area and time) is more important than SLA in determining RGR at cool temperatures. However, such an analysis has not yet been carried out for the below-ground part of plants. Therefore, it is still unclear what the relative contributions are of SRL and RMF to the root length, and whether root absorption activity and root length contribute differently to nutrient acquisition, when plants face low temperature stress. A recent report by Freschet et al. (2015b) suggests that the size ratio of roots to leaves increases as nutrient limitation aggravates, and that RMF contributes more to RLR variation than SRL. It seems that RMF is more important than SRL in determining the root length at cool temperatures, because nutrient limitation can also be aggravated by reducing root-zone temperature. However, despite increased size ratio of roots to leaves, the relative size of roots is generally decreased at cool temperatures (Larigauderie et al., 1991), inferring that RMF and SRL may contribute in different ways to RLR.

Although many studies take temperature as a homogeneous whole, the spatial heterogeneity of temperature (e.g. air and root-zone temperatures) extensively exists either in natural environments (Deanedrummond and Glass, 1983; Walter et al., 2009) or under cultivation conditions (Gosselin and Trudel, 1985; Teitel et al., 1999; Kawasaki et al., 2014). It is well-known that air temperature (*T*_air_) is crucial for plant growth. Over the past decades, an increasing number of studies have shed light on the important role played by root-zone temperature (*T*_root_) in plant growth (Tachibana, 1987; Ahn et al., 1999; Murai-Hatano et al., 2008; Nagel et al., 2009; Poire et al., 2010). However, very few studies have attempted to compare the difference between *T*_air_ and *T*_root_ by independently and separately changing each type of temperature in one experiment. Therefore, it is still unclear how *T*_air_ and *T*_root_ separately affect the relative contributions of various plant components to RGR. Weih and Karlsson (2001) have pointed out that *T*_air_ and *T*_root_ have interactive effects on RGR, N productivity (the rate at which dry matter is produced per unit of N in plant biomass per unit of time) and leaf-N content. It means that plant response to root-zone cooling at optimal *T*_air_ can not be simply predicted as a reverse of response to root-zone warming at low *T*_air_. Thus, it is needed to apply a complete factorial design to distinguish the different roles of *T*_air_ and *T*_root_ in plant growth.

Low temperature can limit resource (e.g. C and nutrients) acquisition by plants. Either *T*_air_ or *T*_root_ limitation alone may lead to unequal accessibilities of above- and below-ground parts to resources. Nevertheless, to maximize growth with minimum resource costs, plants generally tend to balance above- and below-ground resource acquisition capacities to achieve the status of C-nutrient colimitation (Ryser and Eek, 2000; Maire et al., 2013). This mechanism is known as the ‘balanced growth’, ‘optimal partitioning’ or ‘functional equilibrium’ hypothesis (Brouwer, 1963; Davidson, 1969; Shipley and Meziane, 2002), which can be formalized as follows:

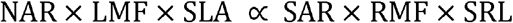

where NAR is plant C net assimilation rate (per unit time and leaf area), and SAR is plant specific nutrient absorption rate (per unit time and root length) (Freschet et al., 2015a). This equation can be further transformed into:

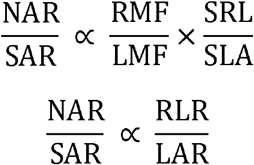

Carbon-nutrient balance may be achieved through various strategies such as maintaining relatively smaller root size but higher SAR, or maintaining relatively constant size/activity ratio of leaves to roots through proportionately increasing NAR and SAR. For instance, in the study of Engels et al. (1992), nutrient uptake by roots is stimulated by increased temperature of maize shoot base (apical shoot meristem and zone of leaf extension) via raising shoot growth, even under low *T*_root_ condition. Getting to know what kind of strategies plants choose can help us better understand the mechanisms of integrated plant responses to temperature limitation.

In this study, we conducted a complete factorial experiment, with two levels of *T*_air_ (high and low) and two levels of *T*_root_ (high and low), to investigate the independent and interactive effects of *T*_air_ and *T*_root_ on leaf and root growth, and carbon and nutrient assimilation in cucumber (*Cucumis sativus* L.) seedlings. The objectives were to examine (1) how *T*_air_ and *T*_root_ affect relative contributions of various plant components to plant growth and resource acquisition, and (2) whether *T*_air_ has different effects on plant carbon-nutrient balance compared to *T*_root_. Cucumber was chosen as a model plant because its growing point locates above the ground surface, which favors separate control of *T*_air_ and *T*_root_, and because it is a major greenhouse crop that is sensitive to low temperature (Terashima et al., 1998) and often grows under different *T*_air_ and *T*_root_ conditions due to artificial control (Gosselin and Trudel, 1985; Teitel et al., 1999; Urrestarazu et al., 2008).

## MATERIAL AND METHODS

### Plant Material and Growth Conditions

Cucumber (*Cucumis sativus* L. cv. Zhongnong No.16) seedlings were hydroponically cultured according to the procedure described by Wang et al. (2016). Briefly, cucumber seeds were pregerminated at 28°C for 26 h, sown onto hydroponic devices and then cultured at 28°C for 30 h under darkness. Germinated seedlings were maintained in hydroponic devices and cultured at 26/18°C (day 10h/night 14h) for 10 days, with 60-80% relative humidity (RH) and approximately 100 μmol photons m^−1^·s^−1^ during the day. Then, seedlings were transplanted onto brown glass bottles placed and cultured at 26/18°C (day 10h/night 14h) for another 5 days, with 60-80% RH and 250 μmol photons m^−2^·s^−1^ during the day. Full-strength Yamazaki nutrient solution (Yamazaki, 1982) at pH 6.0 was used for hydroponics throughout this experiment, and was refreshed every 5 days. The seedlings ready for treatment each had two intact cotyledons, one fully unfolded true leaf and one new leaf beginning to unfold.

### Temperature Treatments

On day 16 after germination, the cucumber seedlings were transferred into temperature controlling devices as described by Wang et al. (2016), which can respectively set and maintain temperature regimes around the shoots and roots. There were two regimes of both *T*_air_ and *T*_root_ in this experiment: 26/18°C (day/night, “high”) and 20/12°C (day/night, “low”) for *T*_air_, and 19±1°C (all-day, “high”) and 13±1°C (constant, “low”) for *T*_root_. A 2 × 2 factorial design was employed to create treatments that included low *T*_air_/low *T*_root_ (L/L), low *T*_air_/high *T*_root_ (L/H), high *T*_air_/low *T*_root_ (H/L) and high *T*_air_/high *T*_root_ (H/H). To assure the comparability of the morphology and biomass allocation of seedlings among different treatments, each treatment lasted until the same stage of seeding development (i.e., for each seedling the second true leaf fully unfolded and the third true leaf was just about to unfold). The actual treatment periods were 10 days for low *T*_air_ treatments (L/L and L/H) and 5 days for high *T*_air_ treatments (H/L and H/H), respectively. Forty seedlings per treatment were cultivated.

### Growth Characteristics

At both the beginning and ending of the treatments, seedlings were harvested to determine growth characteristics (seven replicates, three seedlings each replicate). Fresh leaves and roots were scanned (EPL/HN EXPREL/LION 4990, Japan), and the scanned images were used to quantify leaf area and total root length with WinRHIZO software (LC4800-II LA2400; Sainte-Foy, Canada). Additionally, the path length of each first-order lateral root (LR) on the basal half of main root, and the number of second-order LRs on each first-order LR were quantified with ImageJ software (V1.50b; Abràmoff et al., 2004). The definition of first- and second-order LR was the same as described by Kellermeier et al. (2014). After scanning, fresh plant tissues were separately (root, stem, cotyledon, the first true leaf and the second true leaf) oven-dried at 105°C for 15 min and at 85°C for 48 h, and weighted for the dry mass. Total plant dry mass was calculated as the sum of all plant tissues. For calculating RGR, plants before and after treatment were paired based on the order of total plant dry weight, and then RGR was calculated for paired plants as described by Hunt (1978):

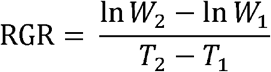

where *W*_1_ and *W*_2_ are the total plant dry mass before and after treatment, respectively, and *T*_2_−*T*_1_ is the treatment period. SLA (cm^2^·g^−1^) and SRL (m·g^−1^) were calculated as the leaf area per leaf dry mass and the root length per root dry mass, respectively. LMF (g·g^−1^) and RMF (g·g^−1^) were estimated as proportions of leaf dry mass and root dry mass of the total plant dry mass, respectively. LAR (cm^2^·g^−1^) and RLR (m·g^−1^) were calculated as the leaf area and root length per total plant dry mass, respectively.

### Element Content and Absorption Rates

After being weighted, the dry tissues of seedlings were ground into fine powder with a mortar and pestle for analysis of element contents. The contents of C and N in seedling tissues were determined by combusting the powder at 900°C within an elemental analyzer (vario PYRO cube, Germany). The contents of P, K, Ca, Mg, Fe, Mn, Zn, and Cu were determined by digesting the powder with nitric acid in a microwave digestion system (MARS 240/50, CEM, USA) and then analyzing with an inductively coupled plasma atomic emission spectrometer (ICP-AES, ICP6300, Britain).

For nutrient elements, the whole plant absorption rate (R_x_, mg element·g^−1^ plant biomass·d^−1^, where x can be N, P, K, Ca, Mg, Fe, Mn, Zn or Cu) and specific absorption rate on a root-length basis (SAR_x_, mg element·m^−1^ root length·d^−1^) were calculated as mean values over the treatment period according to Welbank (1962) as follows:

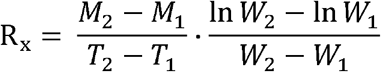

where *M*_1_ and *M*_2_ are the total content of element before and after treatments, respectively, and *R*_L1_ and *R*_L2_ are the total root length before and after treatments, respectively. The estimation of mean 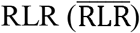 over the treatment period was calculated as dividing R_x_ by SAR_x_.

To compare with N and other elements, the influx of C was also estimated in similar methods, the whole plant net assimilation rates of carbon (R_C_, mg C·g^−1^ plant biomass·d^−1^) and unit leaf rate of carbon (NAR_C_, mg C·cm^−2^ leaf area·d^−1^) were calculated as follows:

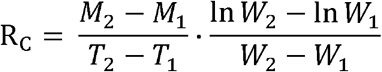

where *L*_A1_ and *L*_A2_ are the total leaf area before and after treatments, respectively. Similarly, the estimation of mean 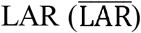 over the treatment period was calculated as dividing R_C_ by NAR_C_.

### Net Assimilation Rate of True Leaves

Gas-exchange was measured on both true leaves with the LI-6400xt gas exchange analyzer (Li-Cor 6400xt, Lincoln, NE, USA) (four seedlings per treatment). Determination started from the third hour of a light period in the last day of treatment. The block temperature was set at the air temperature of the corresponding treatment, and the PAR and air relative humidity were maintained at 1200 μmol·m^−2^·s^−1^ and 60-70%, respectively. Net assimilation rate under 400 μmol CO_2_·mol^−1^ reference CO_2_ concentration (*A*_400_) was recorded.

### Data Analysis

Following the variance partitioning method described by Rees et al. (2010) and Freschet et al. (2015b), we calculated the relative contributions of variance in LMF and SLA to variance in LAR, of RMF and SRL to RLR, of NAR_C_ and 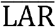 to R_C_, and of SAR_N_ and 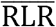 to R_N_. To avoid meaningless results, the variance partitioning was not performed if less than 15% variation was observed in LAR, RLR, R_C_ or R_N_. Instead of relative contributions, direct effects of R_C_ and R_N_ on RGR were worked out using path coefficient analysis (Dewey and Lu, 1959).

The data and the graphs were processed using Microsoft Excel 2016 and Microsoft R Open 3.4.1. For multiple comparisons, data were log_2_ transformed and then subjected to one-way analysis of variance (ANOVA). When ANOVA indicated significant differences (*P* < 0.05), means were compared using Tukey HSD tests (software: IBM SPL/L Statistics 20, IBM Corporation, USA). Two-way ANOVA was performed to compare sources of variation, including *T*_air_, *T*_root_, and the *T*_air_×*T*_root_ interaction.

## RESULTS

### Plant Growth Parameters under Different Temperature Conditions

Seedlings spent less time growing one new leaf under high *T*_air_ conditions (H/L and H/H) than under low *T*_air_ conditions (L/L and L/H). Thus, although elevated *T*_air_ decreased the total dry mass of seedlings at the end of the experiment, it significantly accelerated their RGR (H/L vs L/L, H/H vs L/H; **Table 1**). Elevated *T*_root_ increased total dry mass and RGR at both levels of *T*_air_ (L/H vs L/L, H/H vs H/L). Significant interactive effects of *T*_air_ and *T*_root_ were observed on both total dry mass and RGR. Elevated *T*_air_ significantly decreased leaf area at low *T*_root_ (H/L vs L/L) and total root length at both levels of *T*_root_ (H/L vs L/L, H/H vs L/H). By contrast, elevated *T*_root_ significantly increased total leaf area and total root length at both levels of *T*_air_ (L/H vs L/L, H/H vs H/L). **Table 1** and **Supplementary Figure S1** also display more details about how temperature treatments affected the size of leaves and roots. Compared with older tissues, the 2nd true leaf and the 2nd order LR, which newly developed during treatment, had higher size variation between treatments. Apparently, low *T*_air_ combined with low *T*_root_ resulted in a generally small and slowly-developed L/L seedling. On this basis, elevated *T*_air_ led to a fast-developed but still small seedling (H/L vs L/L), elevated *T*_root_ led to a large but still slowly-developed seedling (L/H vs L/L), and co-elevated *T*_air_ and *T*_root_ led to a large and fast-developed seedling (H/H vs L/L).

**Table 1.** Plant growth parameters of seedlings before and after different temperature treatments. Means with different letters are significantly different (*P* < 0.05, n = 6-7) by Tukey HSD. Source of variation: *F* values and significance (* *P* < 0.05; ** *P* < 0.01; *** *P* < 0.001; ns, not significant) of air temperature (*T*_air_), root-zone temperature (*T*_root_) and *T*_air_× *T*_root_. L/L, low *T*_air_/low *T*_root_; L/H, low *T*_air_/high *T*_root_; H/L, high *T*_air_/low *T*_root_; H/H, high *T*_air_/high *T*_root_.

| Treatment | Total dry mass /mg |  | RGR /mg·g<sup>-1</sup>·d<sup>-1</sup> |  | Leaf dry mass /mg |  | Root dry mass /mg |  | Total leaf area /cm<sup>2</sup> |  | Total root length /m |  |
| --- | --- | --- | --- | --- | --- | --- | --- | --- | --- | --- | --- | --- |
| Before treatment | 183 |  | — |  | 135 |  | 18.9 |  | 51.9 |  | 5.89 |  |
| L/L | 481 | b | 96 | d | 342 | b | 46.5 | b | 95.1 | c | 7.44 | c |
| L/H | 599 | a | 118 | c | 396 | a | 80.1 | a | 139.3 | a | 27.48 | a |
| H/L | 358 | d | 134 | b | 265 | c | 32.0 | d | 92.0 | c | 6.71 | d |
| H/H | 394 | c | 153 | a | 279 | c | 41.7 | c | 116.8 | b | 13.31 | b |
| Source of variation |  |  |  |  |  |  |  |  |  |  |  |  |
| $T_{\text{air}}$ | 409 | *** | 577 | *** | 341 | *** | 369 | *** | 129 | *** | 453 | *** |
| $T_{\text{root}}$ | 79 | *** | 191 | *** | 37 | *** | 228 | *** | 1129 | *** | 2612 | *** |
| $T_{\text{air}} \times T_{\text{root}}$ | 12 | ** | 9 | ** | 8 | ** | 27 | *** | 60 | *** | 254 | *** |
| Treatment | Leaf area /cm<sup>2</sup> |  |  |  |  |  | Root length /cm |  |  |  |  |  |
|  | Cotyledon |  | 1st true leaf |  | 2nd true leaf |  | Main root |  | 1st class LR |  | 2nd class LR |  |
| Before treatment | 18.2 |  | 33.7 |  | — |  | 29.7 |  | 363 |  | 197 |  |
| L/L | 20.2 | a | 42.7 | c | 32.2 | c | 29.9 | b | 391 | c | 323 | c |
| L/H | 19.7 | a | 54.8 | a | 64.9 | a | 40.7 | a | 954 | a | 1754 | a |
| H/L | 19.5 | a | 39.4 | d | 33.1 | c | 29.4 | b | 367 | c | 275 | c |
| H/H | 19.7 | a | 50.8 | b | 46.2 | b | 34.5 | ab | 567 | b | 731 | b |

| Source of variation |  |  |  |  |  |  |  |  |  |  |  |  |
| --- | --- | --- | --- | --- | --- | --- | --- | --- | --- | --- | --- | --- |
| $T_{\text{air}}$ | 0.566 | ns | 31 | *** | 113 | *** | 4.4 | * | 38 | *** | 29 | *** |
| $T_{\text{root}}$ | 0.203 | ns | 330 | *** | 1236 | *** | 27.5 | *** | 187 | *** | 193 | *** |
| $T_{\text{air}} \times T_{\text{root}}$ | 0.848 | ns | 0 | ns | 154 | *** | 3.0 | ns | 21 | *** | 13 | ** |

### Effects of *T*_air_ and *T*_root_ on the Components of Leaf and Root Size

Elevated *T*_air_ significantly raised LAR at each *T*_root_ (**Figure 1**) and RLR at low *T*_root_ (H/L vs L/L), but decreased RLR at high *T*_root_ (H/H vs L/H). Elevated *T*_root_ raised both LAR and RLR at each *T*_air_, and the promoting effect on RLR was obviously stronger at low *T*_air_ than at high *T*_air_. Responses of SLA and SRL to temperature variation were similar to those of LAR and RLR, except for that SRL was not significantly affected by elevated *T*_air_ at high *T*_root_ (H/H vs L/H). As to biomass allocation, elevated *T*_air_ increased LMF and decreased RMF (H/L vs L/L), while elevated *T*_root_ showed reverse trends (L/H vs L/L), leading to unchanged LMF and increased RMF at co-elevated *T*_air_ and *T*_root_ (H/H vs L/L). For different leaves of a seedling, their biomass fractions may respond differently to temperature changes, depending on the leaf order (**Figure 2**). Elevated *T*_air_ significantly increased the allocation ratio of biomass increment of the 2nd true leaf, while elevated *T*_root_ significantly decreased the allocation ratio of biomass increment of cotyledon. For the 1st true leaf, temperature variation had no obvious influence on biomass allocation.

**Figure 1.**
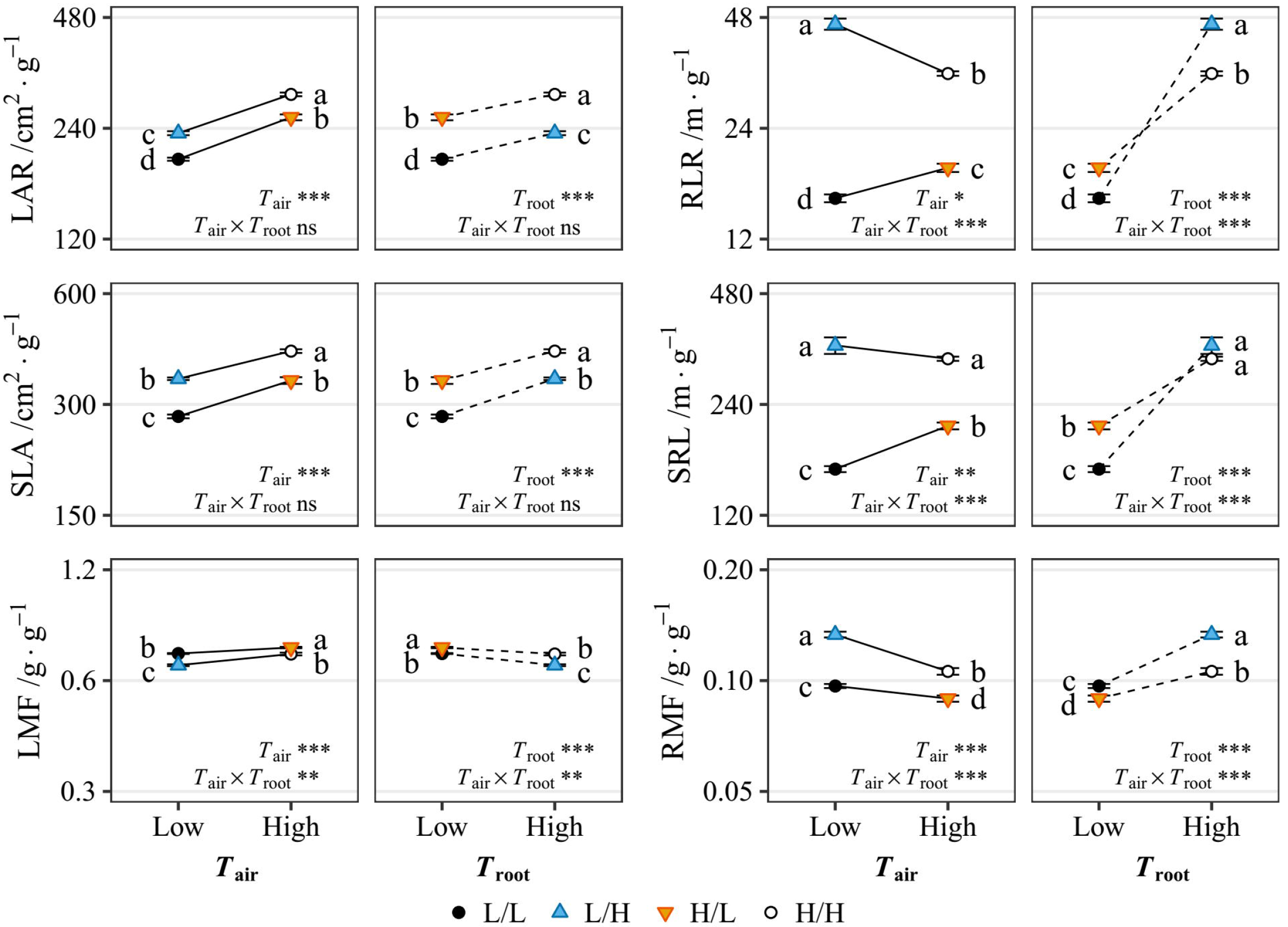
Responses of leaf and root relative size, morphology and biomass allocation to air temperature (*T*_air_) variation (solid lines) and root-zone temperature (*T*_root_) variation (dashed lines). Each variable is expressed on a log_2_-scale. Data points and error bars are means ± standard error (n = 7). Different letters besides each point denote significance at *P* < 0.05 by Tukey’s HSD-test. The significance (*, *P* < 0.05; *** *P* < 0.001; ns, not significant) of interactions between *T*_air_ and *T*_root_ is displayed in each panel. L/L, low *T*_air_/low *T*_root_; L/H, low *T*_air_/high *T*_root_; H/L, high *T*_air_/low *T*_root_; H/H, high *T*_air_/high *T*_root_. The data (in number) of this figure are exhibited in **Supplementary Table S1**.

**Figure 2.**
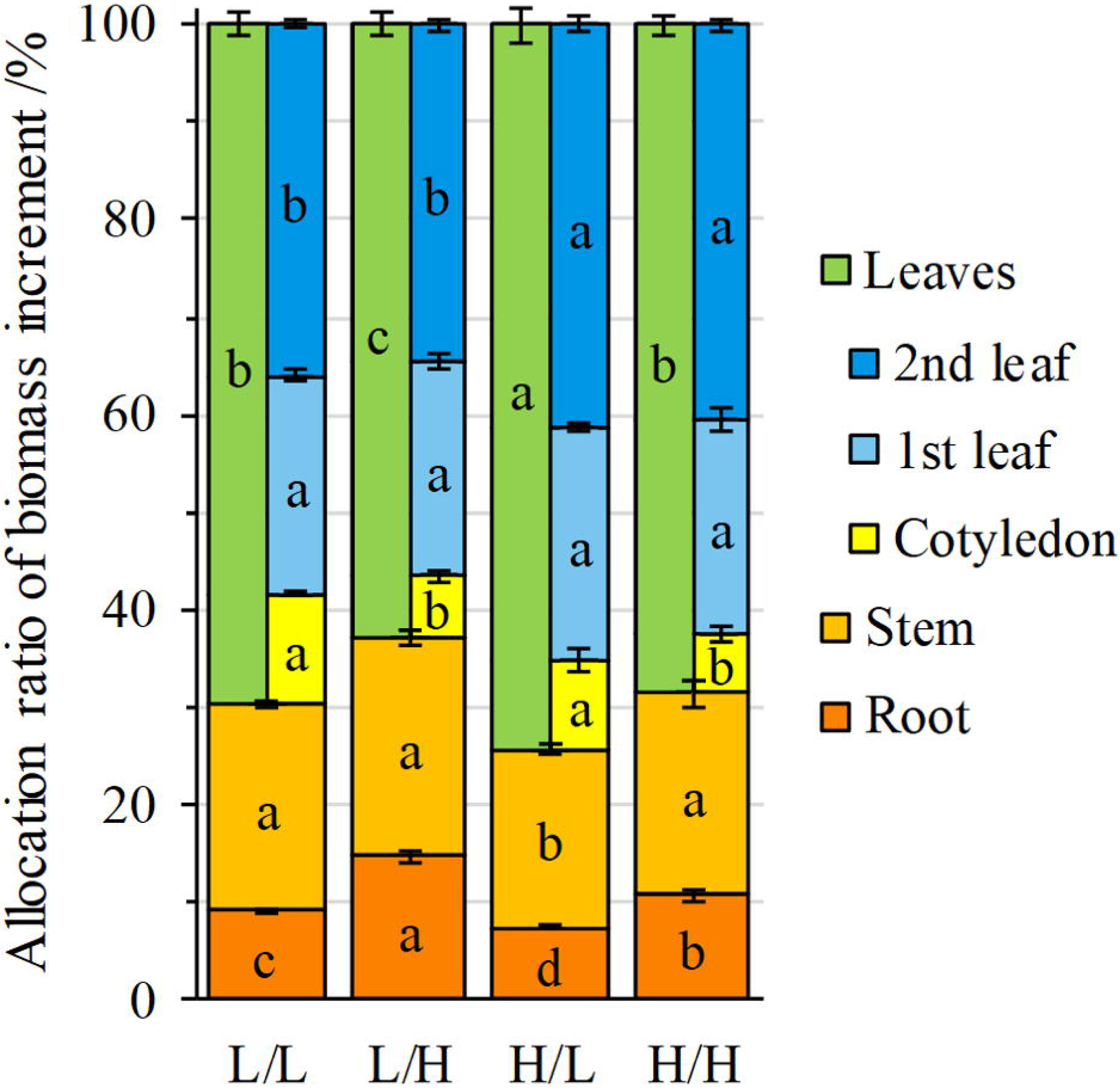
Allocation ratios of biomass increment in different organs during treatment. Boxes and error bars are means ± standard error (n = 7). Different letters besides each point denote significance at *P* < 0.05 by Tukey’s HSD-test. LAR, leaf area ratio; RLR, root length ratio; SLA, specific leaf area; SRL, specific root length; LMF, leaf mass fraction; RMF, root mass fraction. L/L, low *T*_air_/low *T*_root_; L/H, low *T*_air_/high *T*_root_; H/L, high *T*_air_/low *T*_root_; H/H, high *T*_air_/high *T*_root_.

Changes in morphological characteristics (SLA and SRL) generally weighted more than biomass allocation (LMF and RMF) on determining responses of LAR and RLR to temperature variation (**Figure 3**). In the above ground parts, changes in SLA always contributed the major part of the variation in LAR no matter how temperature was changed. The relative contribution of SLA even exceeded 1 since LMF contributed negatively to the total variation in LAR. In the below ground parts, changes in SRL contributed more than RMF to the variation in RLR when *T*_all_ (*T*_air_ + *T*_root_) or *T*_root_ was altered. Specially, only when *T*_air_ changed at high *T*_root_ (L/H vs H/H), the relative contribution of RMF became predominant.

**Figure 3.**
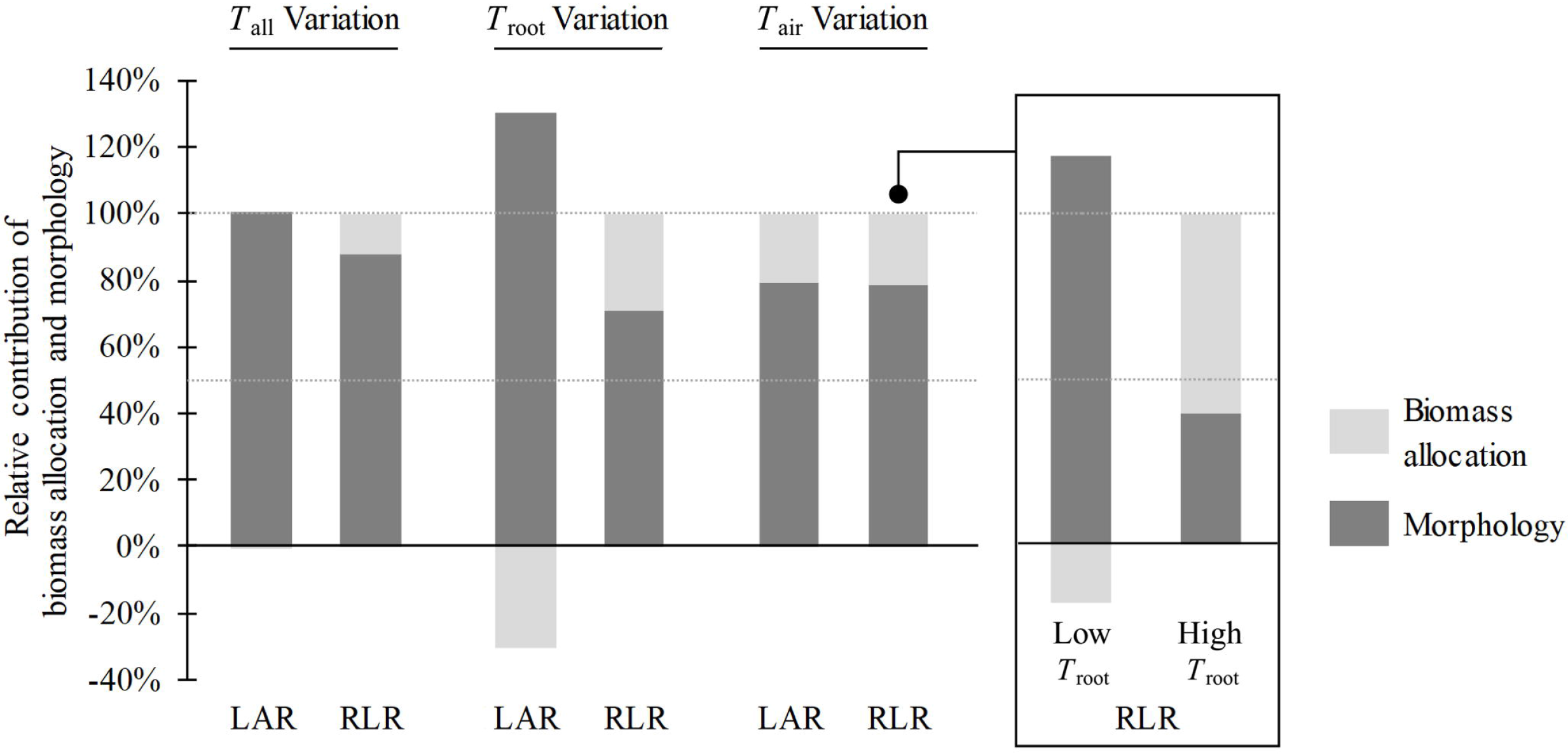
Relative contribution of leaf and root biomass allocation (LMF or RMF) and morphology (SLA or SRL) variables to the total variation in LAR and RLR. The category “*T*_all_ Variation” refers to L/L versus H/H; “*T*_root_ Variation” refers to the mean value of L/L versus L/H and H/L versus H/H; “*T*_air_ Variation” refers to the mean value of L/L versus H/L and L/H versus H/H. Specially, the inserted viewport displays L/L versus H/L (sub-optimal *T*_root_) and L/H versus H/H (optimal *T*_root_) respectively, due to the significant difference between the two conditions. LAR, leaf area ratio; RLR, root length ratio. L/L, low *T*_air_/low *T*_root_; L/H, low *T*_air_/high *T*_root_; H/L, high *T*_air_/low *T*_root_; H/H, high *T*_air_/high *T*_root_.

The ratio of total leaf area to total root length (equivalent to LAR : RLR) varied a lot among different temperature treatments (**Table 2**). Considering the LAR : RLR of H/H seedlings as a balanced standard, *T*_air_ limitation led to a structure with relatively smaller leaves but larger roots in L/H seedlings, while *T*_root_ limitation did the opposite thing on H/L seedlings. Instead of proportionately inhibiting both leaf and root sizes, *T*_all_ limitation led to a high LAR : RLR in L/L seedlings, which was similar to that in H/L seedlings, indicating that root length was more sensitive than leaf area to low temperature. As the components of LAR : RLR, leaf-root morphology (SLA : SRL) and leaf-root biomass allocation (LMF : RMF) varied in a similar way to their product, except for that LMF : RMF was significantly lower in L/L seedlings than in L/H seedlings. This trend was counteracted by SLA : SRL, resulting in no difference in LAR : RLR.

**Table 2.**
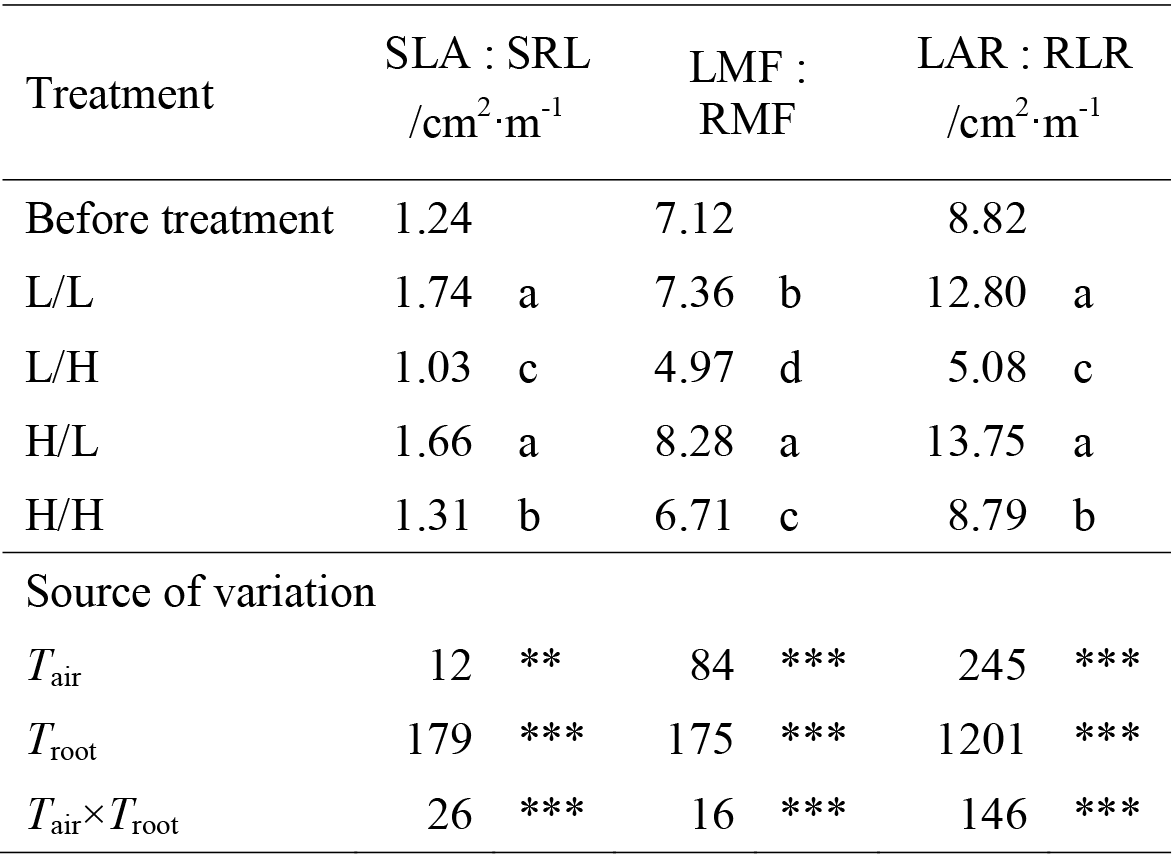
Ratios of leaf : root morphological, biomass fraction and size parameters of seedlings before and after different temperature treatments. Means with different letters are significantly different (*P* < 0.05, n = 7) by Tukey HSD. Source of variation: *F* values and significance (** *P* < 0.01; *** *P* < 0.001) of air temperature (*T*_air_), root-zone temperature (*T*_root_) and *T*_air_×*T*_root_. SLA, specific leaf area; SRL, specific root length; LMF, leaf mass fraction; RMF, root mass fraction; LAR, leaf area ratio; RLR, root length ratio. L/L, low *T*_air_/low *T*_root_; L/H, low *T*_air_/high *T*_root_; H/L, high *T*_air_/low *T*_root_; H/H, high *T*_air_/high *T*_root_.

| Treatment | SLA : SRL<br>/cm<sup>2</sup>·m<sup>-1</sup> |  | LMF :<br>RMF |  | LAR : RLR<br>/cm<sup>2</sup>·m<sup>-1</sup> |  |
| --- | --- | --- | --- | --- | --- | --- |
| Before treatment | 1.24 |  | 7.12 |  | 8.82 |  |
| L/L | 1.74 | a | 7.36 | b | 12.80 | a |
| L/H | 1.03 | c | 4.97 | d | 5.08 | c |
| H/L | 1.66 | a | 8.28 | a | 13.75 | a |
| H/H | 1.31 | b | 6.71 | c | 8.79 | b |
| Source of variation |  |  |  |  |  |  |
| $T_{\text{air}}$ | 12 | ** | 84 | *** | 245 | *** |
| $T_{\text{root}}$ | 179 | *** | 175 | *** | 1201 | *** |
| $T_{\text{air}} \times T_{\text{root}}$ | 26 | *** | 16 | *** | 146 | *** |

### Effects of *T*_air_ and *T*_root_ on Carbon and Nutrient Acquisition and Allocation

At each *T*_root_, elevated *T*_air_ significantly raised both R_C_ and R_N_, while elevated *T*_root_ only raised R_N_ (**Figure 4**). As to specific resource acquiring rates, elevated *T*_air_ raised both NAR_C_ and SAR_N_, and the promoting effect on SAR_N_ was stronger at high *T*_root_ than at low *T*_root_. Elevated *T*_root_ had no significant influence on both NAR_C_ and SAR_N_ at low *T*_air_, and had a negative effect on NAR_C_ but a positive effect on SAR_N_ at high *T*_air_. The response of photosynthetic capacity to temperature variation was different from that of NAR_C_. All elevated-temperature treatments significantly increased the *A*_400_ (net photosynthetic rate) in both true leaves of seedlings (compere H/L, L/H and H/H versus L/L; **Table 4**). However, compared to H/H seedlings, *T*_root_ limitation did not affect the *A*_400_ in any leaf of H/L seedlings, and *T*_air_ limitation only decreased the *A*_400_ in the 2nd true leaf of L/H seedlings. No significant interaction between *T*_air_ and *T*_root_ was observed in R_C_, R_N_ and NAR_C_ (**Figure 4**).

**Figure 4.**
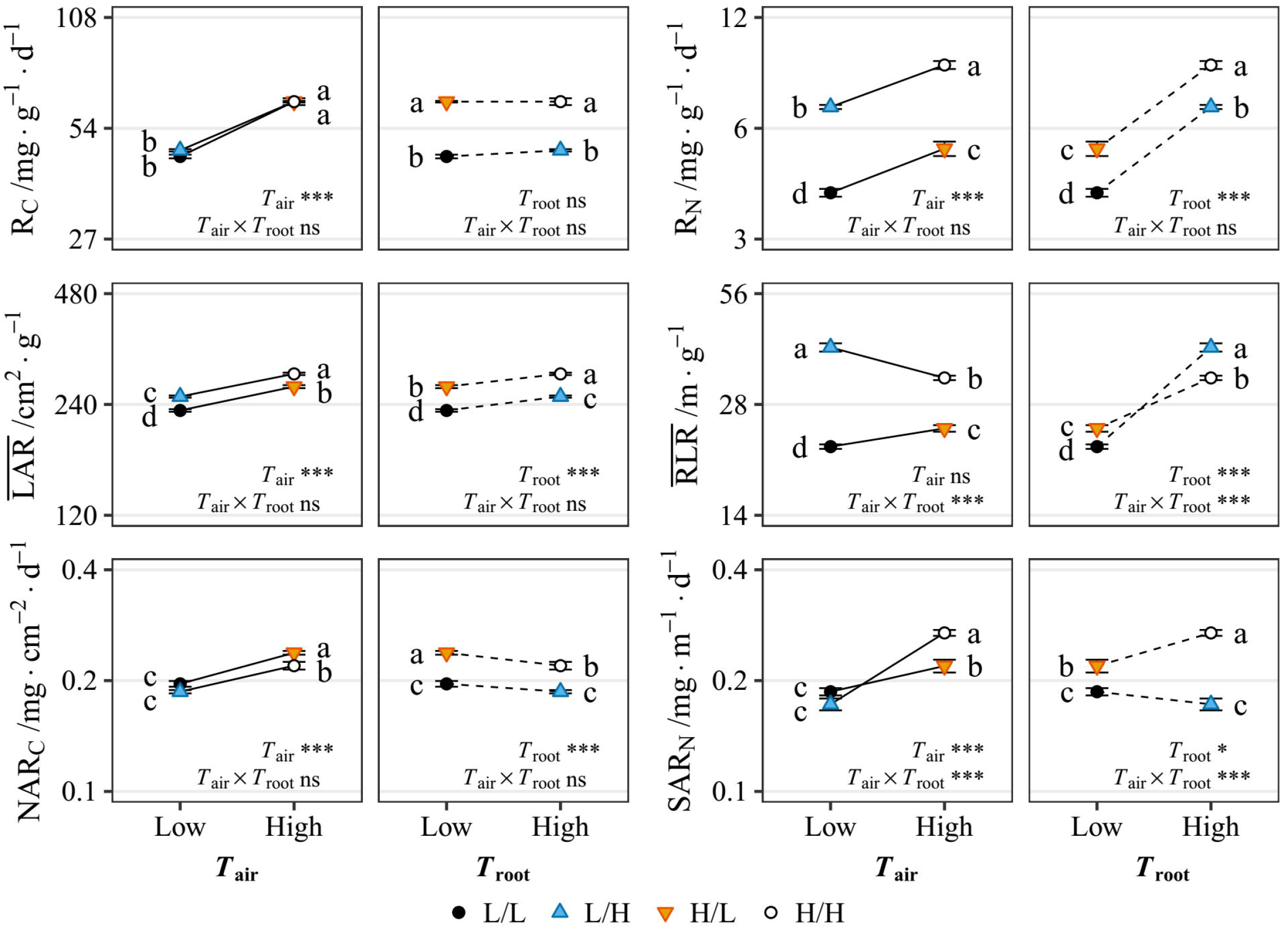
Responses of leaf and root size and assimilation/absorption rate to air temperature (*T*_air_) variation (solid lines) and root-zone temperature (*T*_root_) variation (dashed lines). Each variable is expressed on a log_2_-scale. Data points and error bars are means ± standard error (n = 7). Different letters besides each point denote significance at *P* < 0.05 by Tukey’s HSD-test. The significance (*, *P* < 0.05; ** *P* < 0.01; *** *P* < 0.001; ns, not significant) of interactions between *T*_air_ and *T*_root_ is displayed in each panel. L/L, low *T*_air_/low *T*_root_; L/H, low *T*_air_/high *T*_root_; H/L, high *T*_air_/low *T*_root_; H/H, high *T*_air_/high *T*_root_. The data (in number) of this figure are exhibited in **Supplementary Table S2**.

When *T*_all_ was changed, NAR_C_ contributed 38% of the variation in R_c_, and SAR_N_ and 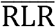 contributed almost equally to the variation in R_N_ (**Figure 5**). When *T*_root_ was changed, 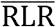 contributed a major part of the variation in R_N_. When *T*_air_ was changed, NAR_C_ contributed 57% of the variation in R_C_, and SAR_N_ contributed a major part of the variation in R_N_.

**Figure 5.**
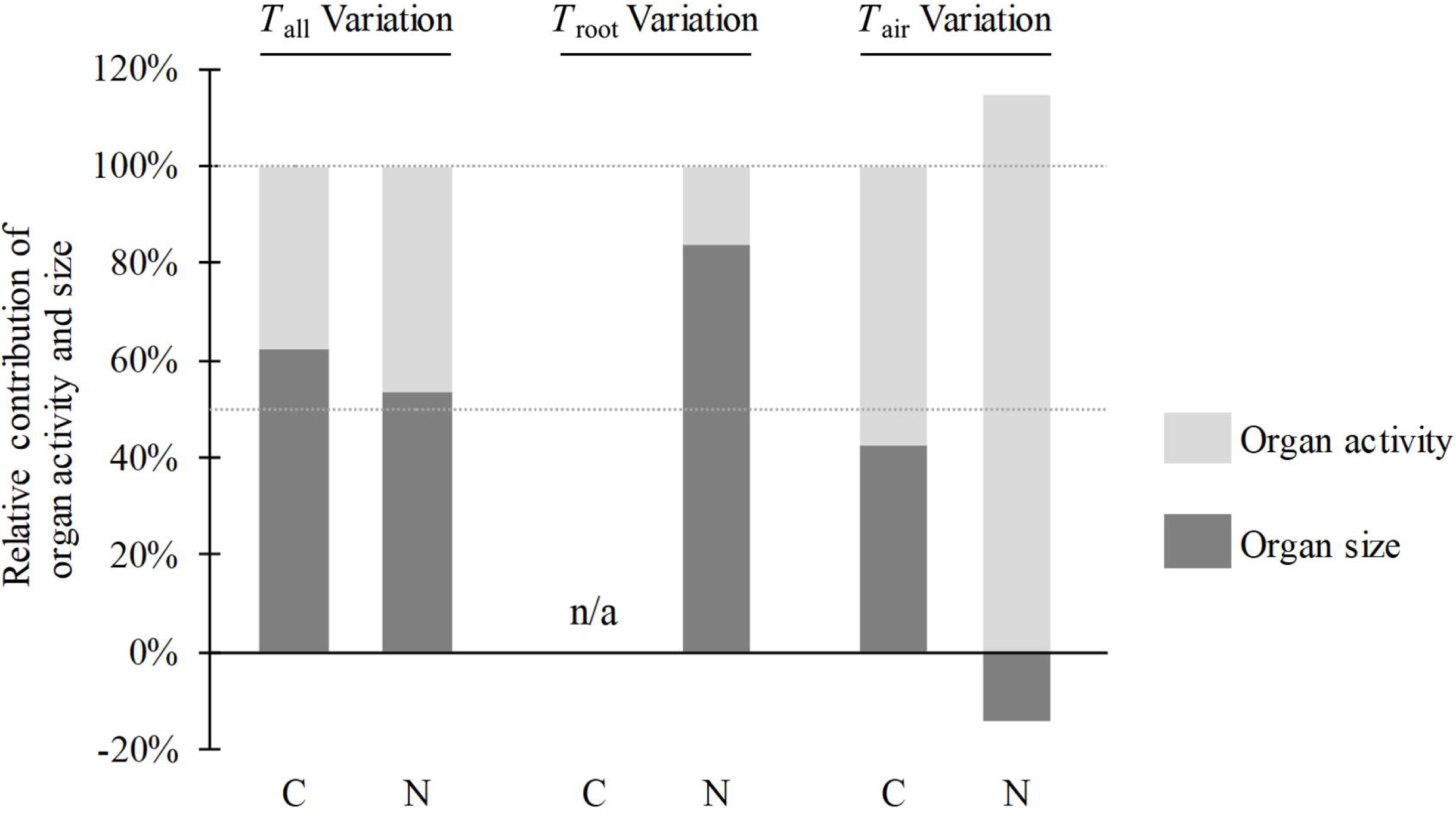
Relative contribution of leaf and root assimilation/absorption activity (NAR_C_ or SAR_N_) and size (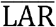 or 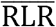) variables to the total variation in R_C_ and R_N_. The category “ *T*_all_ Variation” refers to L/L versus H/H; “*T*_root_ Variation” refers to the mean value of L/L versus L/H and H/L versus H/H; “*T*_air_ Variation” refers to the mean value of L/L versus H/L and L/H versus H/H. n/a, not applicable, as no more than 15% variation in R_C_ was observed when changing *T*_root_. L/L, low *T*_air_/low *T*_root_; L/H, low *T*_air_/high *T*_root_; H/L, high *T*_air_/low *T*_root_; H/H, high *T*_air_/high *T*_root_.

The value of R_c_ : R_N_ is equal to the ratio of newly gained total carbon to nitrogen per day. Considering the Rc : RN of H/H seedlings as a balanced standard, *T*_air_ limitation did not influence the ratio in L/H seedlings. This was mainly because of the counteracting effect of decreased root absorption activity (increased NAR_C_ : SAR_N_) and increased root size (decreased 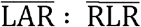) (**Table 3**). *T*_root_ limitation raised R_C_ : R_N_ in H/L seedlings by increasing both NAR_C_ : SAR_N_ and 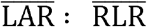. *T*_all_ limitation led to aggregated nitrogen limitation in L/L seedlings, similar to that in H/L seedlings. This similarity was due to no significant difference in either NAR_C_ : SAR_N_ or 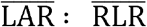 between L/L and H/L seedlings.

**Table 3.** Ratios of leaf : root average size and resource acquiring capacity, and relative carbon : nitrogen accumulation rate of seedlings during different temperature treatments. Means with different letters are significantly different (*P* < 0.05, n = 7) by Tukey HSD. Source of variation: *F* values and significance (** *P* < 0.01; *** *P* < 0.001; ns, not significant) of air temperature (*T*_air_), root-zone temperature (*T*_root_) and *T*_air_×*T*_root_. L/L, low *T*_air_/low *T*_root_; L/H, low *T*_air_/high *T*_root_; H/L, high *T*_air_/low *T*_root_; H/H, high *T*_air_/high *T*_root_.

| Treatment | $\overline{\text{LAR}} : \overline{\text{RLR}}$,<br>/cm<sup>2</sup>·m<sup>-1</sup> | | NAR<sub>C</sub> :<br>SAR<sub>N</sub> | | R<sub>C</sub> : R<sub>N</sub> | |
| --- | --- | --- | --- | --- | --- | --- |
| L/L | 10.76 | a | 1.05 | a | 11.33 | a |
| L/H | 6.32 | c | 1.09 | a | 6.87 | b |
| H/L | 11.15 | a | 1.10 | a | 12.23 | a |
| H/H | 8.80 | b | 0.82 | b | 7.17 | b |

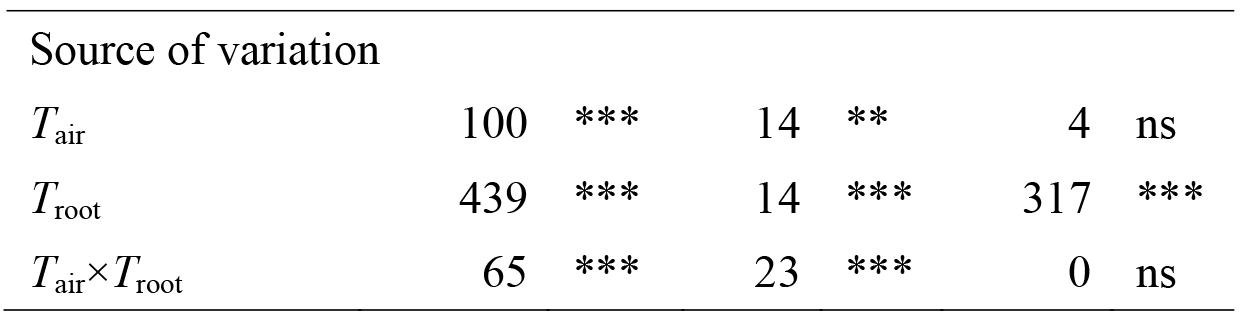

The allocation of newly gained carbon and nitrogen to each part of a seedling was not always proportional to that of biomass, and was distinct among different organs (Figure 6A and B, **Figure 2**). Nitrogen allocation was apparently more flexible than carbon allocation. Newly gained nitrogen was allocated more to new leaf, stem and root than to old leaves. Such trend of heterogeneity was more apparent at high *T*_air_ (H/H and H/L). As a result of aggregated nitrogen limitation, L/L and H/L seedlings had relatively higher carbon concentration and lower nitrogen concentration, and thus higher C/N ratios than L/H and H/H seedlings (Figure 6C, D and E). However, the more heterogeneous nitrogen allocation decreased the C/N ratios of the second leaf, stem and root (compare H/L versus L/L, or compare H/H versus L/H).

**Figure 6.**
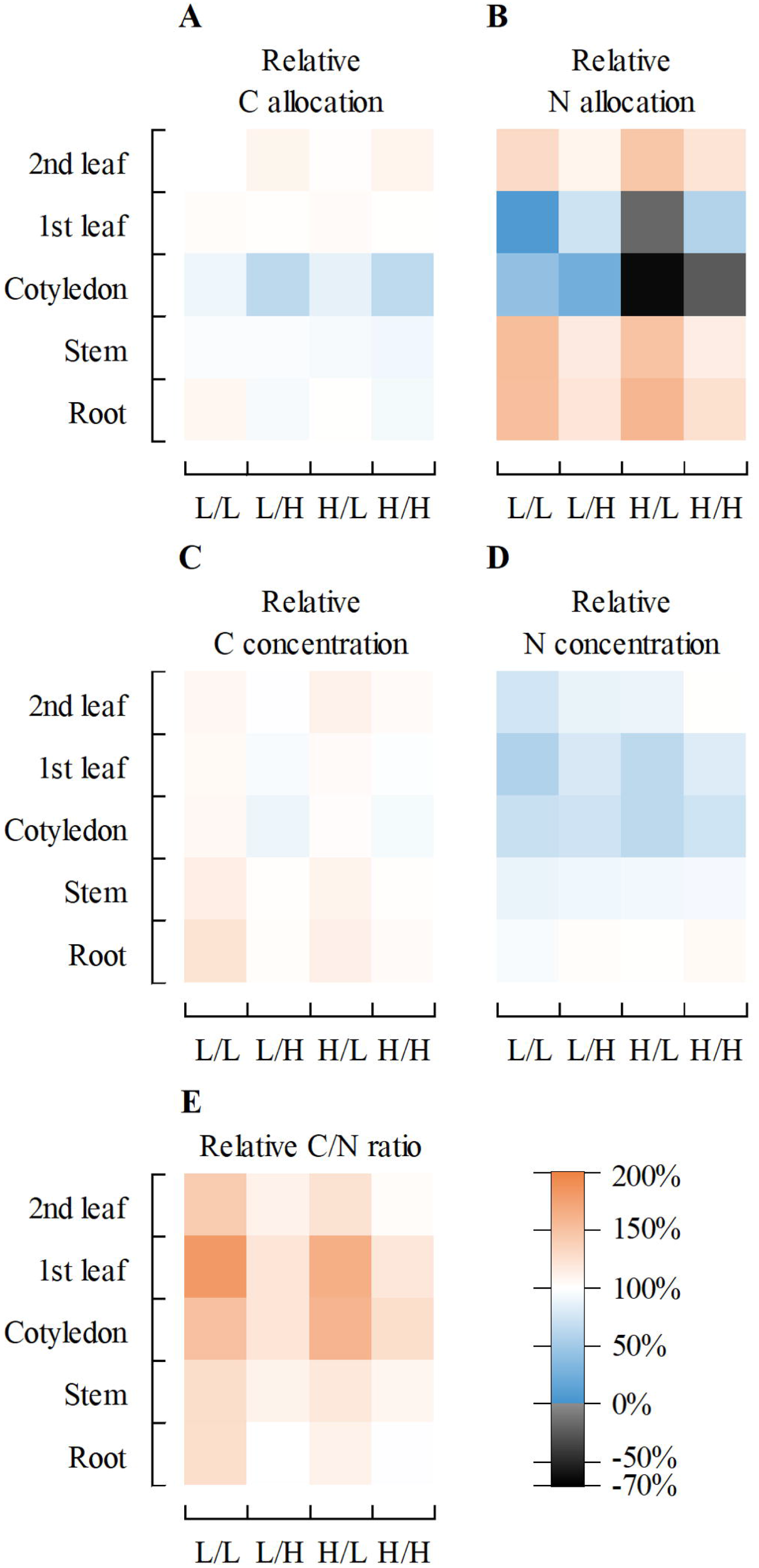
Relative allocation of carbon (A) and nitrogen (B), relative concentration of carbon (C) and nitrogen (D), and relative C/N ratio (E) in each part of the treated seedlings. (A)(B) relative allocation was calculated through dividing the ratio of carbon or nitrogen allocated to each part by the ratio of biomass allocated to correspond part. (C)(D) (E) relative values was calculated through dividing the value in each part by those in the seedlings before treatment. Specially, values in the second true leaves were divided by those in the first true leaves of un-treated seedlings. Negative value of relative allocation indicates net efflux rather than influx of element in the correspond part of plant during treatment. L/L, low *T*_air_/low *T*_root_; L/H, low *T*_air_/high *T*_root_; H/L, high *T*_air_/low *T*_root_; H/H, high *T*_air_/high *T*_root_. The data (in number) of this figure and biomass allocation are exhibited in **Supplementary Table S3**.

The final networks under various conditions were illustrated in **Figure 7**. As the components of RGR, R_C_ and R_N_ had similar direct effects on determining RGR when *T*_all_ was changed (**Figure 7A**). R_C_ had a higher direct effect when *T*_air_ was changed (**Figure 7C, E**), and R_N_ had a higher direct effect when *T*_root_ was changed (**Figure 7B, D**).

**Figure 7.**
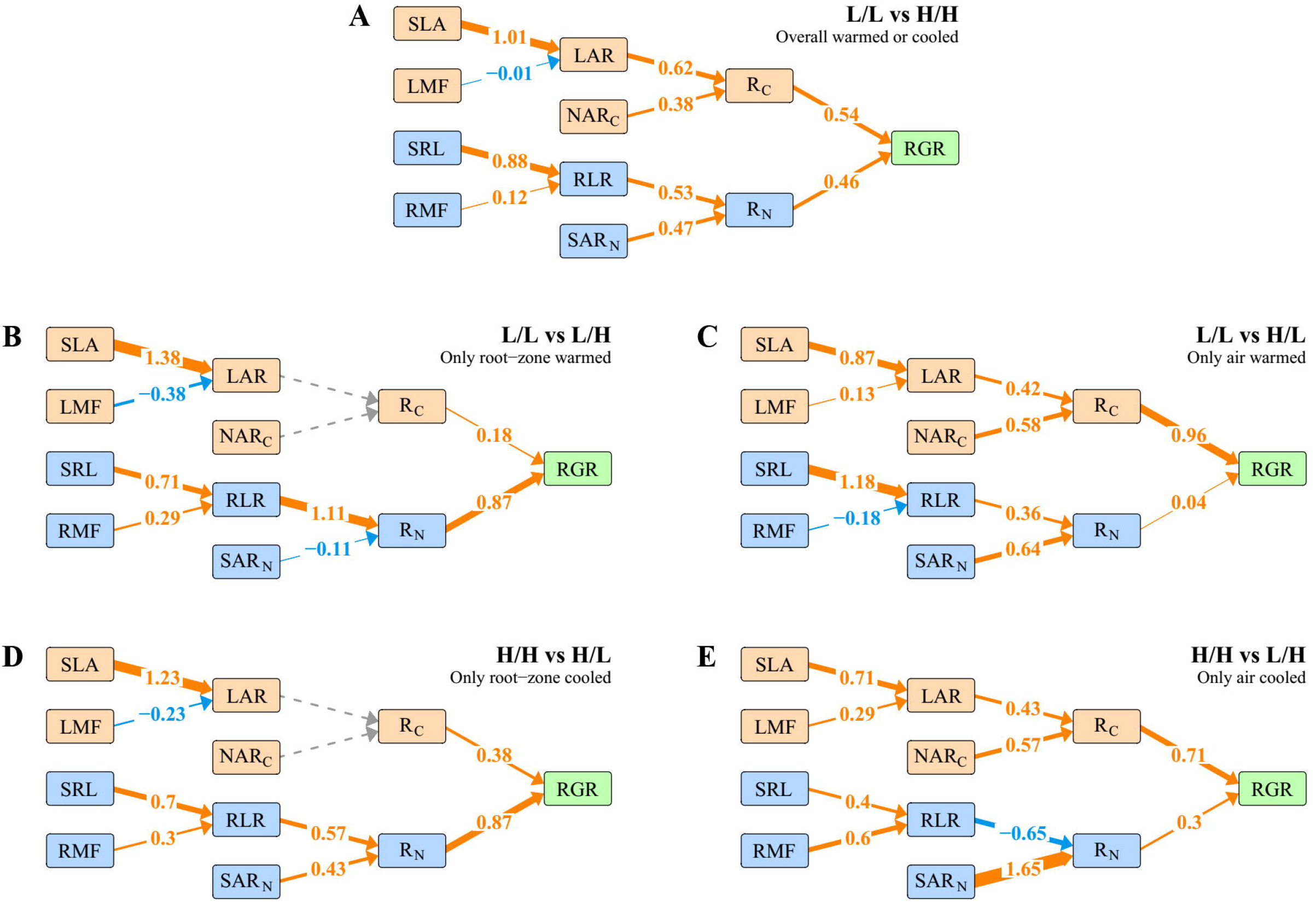
Networks of relative contribution among leaf and root morphology, biomass allocation, size and capacity variables, and direct path coefficients of carbon and nitrogen acquisition rate to relative growth rate under different conditions of temperature variation. (A) L/L vs H/H seedlings, overall warmed or cooled; (B) L/L vs L/H seedlings, only root-zone was warmed; (C) L/L vs H/L seedlings, only air was warmed; (D) H/H vs H/L seedlings, only root-zone was cooled; (E) H/H vs L/H seedlings, only air was cooled. L/L, low *T*_air_/low *T*_root_; L/H, low *T*_air_/high *T*_root_; H/L, high *T*_air_/low *T*_root_; H/H, high *T*_air_/high *T*_root_.

## DISCUSSION

### Morphology Responds More than Mass Allocation to Temperature Variation in Both Leaf and Root

In this study, changes in SLA always accounted for a major part of the temperature-induced variation in LAR, irrespective of *T*_all_, *T*_air_ or *T*_root_ (**Figure 3**). Similar results were reported by Loveys et al. (2002) and Poorter et al. (2012), where temperature was managed based on *T*_all_ or *T*_air_. However, very few studies to date has attempted to examine the *T*_root_-induced variation in SLA. According to the results reported by Weih and Karlsson (2001) and Danyagri and Dang (2014), the variation of SLA is more consistent with that of LAR when compared to LMF, indicating that SLA may contribute more to LAR.

For the below-ground parts, SRL contributed more than RMF to the temperature-induced variation in RLR, irrespective of *T*_all_, *T*_air_ or *T*_root_ (**Figure 3**). Similar trends for *T*_root_ could be obtained based on the data reported by Tachibana (1982) and Engels et al. (1992). However, this trend is opposite to root response to nutrient regulation reported by Freschet et al. (2015b), that is, RMF contributed more than SRL to the nutrient-induced variation in RLR. One possible reason for the reverse trends is that temperature variation also has significant effects on the hydraulic status of roots (Lee et al., 2004; Lee et al., 2005), which plays an important role in determining root morphology (Wan et al., 1999). Another reason is probably due to the difference in culture medium. It seems that, probably because hydroponics favors root elongation better than soils or sands due to lower mechanical impedance (Bengough and Mullins, 1990), the contribtuion of SRL to RLR was predominent under our hydroponic conditions. However, Engels et al. (1992) found that SRL predominated in RLR under both hydroponic and soil culture conditions, indicating the crucial role played by SRL in determining RLR.

### The Role of Root Activity and Size in Nitrogen Uptake Depends on Temperature Management Strategies

When *T*_air_ was elevated alone, NAR_C_ contributed more than LAR to total C assimilation (**Figures 4 and 5**). However, the contribution of NAR_C_ was reduced when *T*_all_ was elevated and was nearly eliminated when *T*_root_ was elevated alone, indicating the inhibiting role of elevated *T*_root_ in NAR_C_. NAR_C_ is the result of leaf photosynthetic rate minus total plant respiration per unit leaf area. Since both leaf photosynthetic capacity and LAR increased at elevated *T*_root_ (the higher LAR, the lower leaf mass per unit area; **Table 4** and **Figure 1**), stimulated respiration rate should be mainly responsible for the reduction in NAR_C_ at elevated *T*_root_. Additionally, because the maintenance respiration rate is generally higher in roots than in leaves (Lambers et al., 1983), and because root respiration increase with increasing temperature (Atkin et al., 2005), the increased RMF : LMF ratio also contributed to the reduction of net carbon accumulation at elevated *T*_root_ in this study. For nitrogen absorption, compared with *T*_root_, *T*_air_ had only a slight influence on root size but a predominant effect on SAR_N_ (**Figure 4**). Such distinct impacts resulted in that when *T*_air_ was changed, SAR_N_ contributed more than 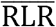 to the variation of R_N_, but this trend was reversed when *T*_root_ was changed (**Figure 5** and Figure 7). It is generally comprehensible that *T*_air_ has less influence on root size than *T*_root_, since the *T*_root_ has more direct effects on root hydraulic status (Wan et al., 1999; Lee et al., 2004; Lee et al., 2005). Supportive results can be found in the researches of Engels and Marschner (1990) and Larigauderie et al. (1991), although in these studies data were not presented in the form of RLR or root area ratio. The influence of *T*_air_ on nutrient uptake is generally considered to be regulated by sugar signals (Stitt and Krapp, 1999). For instance, exogenous application of sugars can increase nitrate reductase activity (Reda, 2015). Moreover, a bZIP transcription factor *Arabidopsis* ELONGATED HYPOCOTYL5 (HY5) has be reported to mediate sucrose signal and promote root nitrate uptake by activating *NRT2.1* (Cerezo et al., 2001; Chen et al., 2016). A previous study (Engels et al. 1992) has reported a promotion effect of elevated shoot base temperature on nutrient translocation rates per unit root fresh weight, but this effect was not examined at elevated *T*_root_. Our observation indicates that the sink-mediated regulation, induced by *T*_air_ management, may overtake the direct effects of *T*_root_ on length-based SAR_N_. However, Weih and Karlsson (2001) showed that raising *T*_root_ was more efficient than raising *T*_air_ in increasing nitrogen uptake rate per unit root dry weight. Actually, this result doesn’t go against our observation, since the trend was also reversed when transforming length-based SAR_N_ into weight-based SAR_N_ (**Supplementary Table S2**) by multiplying SRL in our study. Thus, the significant influence of *T*_root_ on SRL may be partly responsible for counteracting the extent of *T*_root_ effect on length-based SAR_N_.

**Table 4.**
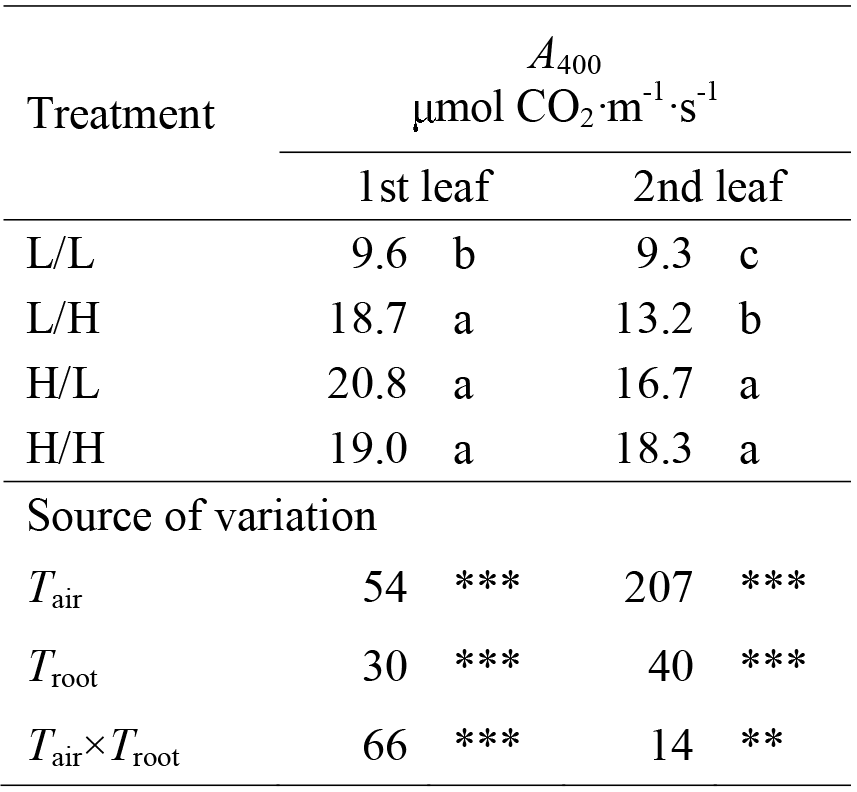
Net photosynthetic rates of cucumber true leaves at 400 μmol CO_2_·mol^−1^ reference CO_2_ concentration. Means with different letters are significantly different (*P* < 0.05, n = 7) by Tukey HSD. Source of variation: *F* values and significance (** *P* < 0.01; *** *P* < 0.001) of air temperature (*T*_air_), root-zone temperature (*T*_root_) and *T*_air_×*T*_root_. L/L, low *T*_air_/low *T*_root_; L/H, low *T*_air_/high *T*_root_; H/L, high *T*_air_/low *T*_root_; H/H, high *T*_air_/high *T*_root_.

R_C_ and R_N_ are additive in affecting RGR, and both are predominant. When *T*_air_ was elevated, R_C_ and RN were proportionally promoted. However, R_C_ accounted for more variation in RGR than R_N_. This is because the concentration of carbon is higher than that of nitrogen in plant tissue. In contrast to the situation at *T*_air_, when *T*_root_ was elevated, R_C_ was not affected, and R_N_ (together with other elements absorbed by roots) became the main reason for variation in RGR (**Figure 4 and Figure 7**). The trends mentioned above were not applicable to the situation at elevated *T*_all_.

### Adaptive Phenotypic Plasticity in Response to Altered *T*_air_ and *T*_root_ in Plants

In a heterogeneous temperature (*T*_air_ vs *T*_root_) environment, cucumber seedlings tended to invest less biomass and generate relatively smaller organ in the cooler zone. Such passive response would potentially strengthen rather than relieve the limitation of the corresponding resource. This trend seems to go against the functional equilibrium hypothesis, that a plant would invest more biomass in the organ responsible for acquiring the most limiting resource (Brouwer, 1963; Freschet et al., 2015b). Based on previous studies regarding biomass allocation, there are both supportive (Tachibana, 1982; Clarkson et al., 1986; Delucia et al., 1992; Danyagri and Dang, 2014) and opposed (Davidson, 1969; Engels and Marschner, 1990; Li et al., 1994; Yan et al., 2012) evidences. It is possible that various factors (e.g. species, temperature, growth medium and ontogenetic stage) may also have impacts on the direction of biomass allocation. For the morphological response, low temperature-induced hydraulic limitation (Murai-Hatano et al., 2008; Wang et al., 2016) and abscisic acid accumulation (Zhang et al., 2008; Ntatsi et al., 2014), which are generally less combined with nutrient and/or light limitation, are both responsible for retarding leaf and root growth (Walter et al., 2009; Pantin et al., 2011). Therefore, in response to temperature variation, changes in the trend of biomass allocation and relative leaf : root size ratio may be more passive rather than adaptive compared with those in respond to nutrient and/or light variation.

According to the above response and the ‘balanced growth’ hypothesis, NAR_C_ : SAR_N_ should vary against the trend of LAR : RLR in order to maintain balanced carbon-nutrient acquisition (i.e. R_C_ : R_N_). Actually, in this study, NAR_C_ : SAR_N_ increased no matter which of *T*_air_ and *T*_root_ was cooled. At low *T*_air_, the higher NAR_C_ : SAR_N_ counterbalanced the lower LAR : RLR, resulting in a relatively constant R_C_ : R_N_. At low *T*_root_, however, the higher NAR_C_ : SAR_N_ was accompanied by a higher LAR : RLR, leading to a large increase in R_C_ : R_N_ (i.e. carbon accumulation or nitrogen limitation). As discussed above, the counteracting effect of root respiration on total carbon acquisition could be one of the main reasons for carbon accumulation when *T*_root_ was cooled. In addition, the sugar-induced increase in nitrogen acquisition could be largely inhibited by limited transporter activity (Reay et al., 1999) and root size at low *T*_root_. In response to nitrogen limitation, newly gained nitrogen was more unevenly distributed between older leaves and other organs to ensure adequate nitrogen concentration for growth in the latter (L/L vs L/H, H/L vs H/H seedlings, **Figure 6**).

### The *T*_air_ and *T*_root_ Interactively Determine Root Size

In this study, the *T*_air_ and *T*_root_ interactively affected RGR, and all root length- and root biomass-related parameters. The interaction effects on RGR were also reported in Larigauderie et al. (1991) and Weih and Karlsson (2001), which showed that increasing *T*_air_ or *T*_root_ alone had a greater promotion on RGR than increasing *T*_all_. Interestingly, interaction effects were not observed in R_C_ and R_N_, neither in LAR and NAR_C_, all of which are components of RGR. Thus the only possible interpretation is that elevated *T*_root_ had a weaker effect on uptake of other elements except for nitrogen at high *T*_air_ than at low *T*_air_. In addition, elevated *T*_root_ had a weaker effect on LR length, particularly the length of the second order LRs, at high *T*_air_ than at low *T*_air_ (**Table 1** and **Supplementary Figure S1**), since the treatment period for seedlings was shorter at high *T*_air_ (thus less accumulated *T*_root_) (KASPAR and BLAND, 1992). This infer that the *T*_air_ and *T*_root_ interaction effects on root length and root biomass might be further reduced by the initiation and development of lateral roots. Generally, the second order LRs were much thinner than main roots and the first order LRs (about 0.2mm vs 0.5~2mm), indicating that elevated *T*_root_ led to a much higher SRL at low *T*_air_ than at high *T*_air_. This interaction effect can interpret some exceptions in the observed trends, e.g., contributed less than RMF to RLR when *T*_air_ was changed at high *T*_root_ (**Figure 3**), and that *T*_air_ variation had less influence on LAR : RLR, SLA : SRL and LMF : RMF at low *T*_root_ than at high *T*_root_ (**Table 2**).

In this study, cucumber seedlings with the same number of leaves were compared after different treatments, and this was originally designed to avoid ontogenetic effects. The period used for new leaf initiation was changed only by varying *T*_air_. This is consistent with the report of Savvides et al. (2016), which showed that the rate of cucumber leaf initiation was completely determined by apical bud temperature independent of the temperature of other plant organs. Although apical bud temperature was not monitored in our experiment, it can be regarded as varying along with *T*_air_ rather than *T*_root_. Field experiment on tomato also reported that cropping was delayed by low *T*_air_ irrespective of *T*_root_ (Jones et al., 1978). However, the treatment period aiming for a uniform shoot developmental stage induced ontogenetic drift in roots, as discussed above. Thus, besides plant growth, the distinct influence of *T*_air_ and *T*_root_ variation on shoot and root development or phenology should also be taken into consideration when designing temperature control strategy for experiment or for protected cultivation.

## CONCLUSION

Our results revealed the distinct effects of *T*_air_ and *T*_root_ on cucumber seedling growth. The primary influence of cooling *T*_root_ on seedling growth was decrease in SRL, which was the main contributor to decrease in RLR. Lower RLR contributed the major part of decrease in total nitrogen acquisition, which finally retarded RGR in seedlings at lower *T*_root_. Variation in *T*_root_ didn’t affect net carbon fixation, although cooling *T*_root_ also decreased LAR mainly via reducing SLA. The major effect of decreasing *T*_air_ on seedling growth was decrease in the capacities of carbon assimilation in leaves and nitrogen absorption by roots, which contributed more than LAR and RLR to the reduction in total resource acquisition. The ratio of carbon : nitrogen acquisition was maintained at a relatively constant level when *T*_air_ was changed, but was increased by decreasing *T*_root_. The interactive effect of *T*_air_ and *T*_root_ was mainly observed on RGR and root growth related variables.

## AUTHOR CONTRIBUTIONS

Conceived and designed the experiments: XW, YT and LG. Performed the experiments: XW. Analysed the data: XW and YT. Wrote the paper: XW and YT.

## FUNDING

This work was supported by the Special Fund for the National Key Research Development Program of China (2016YED201003) and the earmarked fund for the China Agriculture Research System (CARS-25-C-12).

